# Contrasts and similarities in the transcriptomic response to antimicrobial coinage metals in *Escherichia coli*

**DOI:** 10.1101/2025.08.12.669831

**Authors:** Daniel A. Salazar-Alemán, Ashley McGibbon, Raymond J. Turner

**Affiliations:** Department of Biological Sciences, University of Calgary, Calgary, Alberta, Canada

**Author notes:** Address correspondence to Raymond J. Turner. Ashley McGibbon, Department of Microbiology and Immunology, McGill University, Montreal, Quebec, Canada.

## Abstract

With the rise of resistance to last resort antibiotics, metal-based antimicrobials have re-emerged as an alternative to prevent and manage infections. The group 11 metals (copper, silver, gold), historically known for their usage in coins and similar chemical properties, have demonstrated promising bactericidal activity. Despite their efficiency, we do not have a complete understanding for how bacteria are eradicated by metal ions and how they respond to metal-induced stress. One understudied aspect of these metal-bacteria interactions are prolonged exposure models, as other studies tend to focus on the acute toxic response of antimicrobial metals. We used RNA-seq profiling to understand the *Escherichia coli* physiological response to sublethal inhibitory antimicrobial coinage metal stress after 10 hours of incubation. Gene expression patterns of the adaptive and intrinsic response elicited by each metal were identified, including increased essential metal uptake (Ag, Cu, Au), cysteine biosynthesis (Cu, Au), change of the metal ion oxidation state (Cu, Au), efflux of metal stressor (Cu), protein translation and ribosome biogenesis (Au), and cell envelope stress response (Ag). In this paper, we highlight the remarkable differences and similarities in the transcriptomic response profile of *E. coli* to these antimicrobial metal elements.

**IMPORTANCE:** Dogma has existed that all antimicrobial metals kill bacteria the same way, leading to the assumption that bacteria respond the same way to metal toxicity. Nowadays, we have a better understanding why some metal elements are more toxic than others, but questions remain in relation to how bacteria adapt to survive and thrive when challenged by different metal-based antimicrobials. Our study advances the field by characterizing the type of bacterial response(s) to acclimate and grow during a prolonged exposure of silver, copper and gold – metallic elements that are known for their antimicrobial activity. Taking advantage of well-characterized *Escherichia coli*, we propose a model that summarizes our findings after comparing the shared and unique responses to each of these metals. This information enhances our understanding of bacterial tolerance to metal-based antimicrobials, which can lead to improved drug development strategies as society continues to search for alternatives against antibiotic-resistant pathogens.

## INTRODUCTION

Metal-based antimicrobials (MBAs) have been used for millennia. Ancient civilizations, such as the Persians and Phoenicians, used silver and copper in vessels to prevent water fouling (1). British settlers in America preserved water by suspending precious metal coins into wooden casks (2). The “father of bacteriology” Robert Koch explored the effectiveness of silver and cyanide-gold compounds against *Mycobacterium tuberculosis* through the last two decades of the 19^th^ century (3). Antimicrobial approaches that harnessed the biocidal properties of metals enjoyed the spotlight until Sir Alexander Fleming’s discovery of penicillin in 1928 (4), which subsequently caused antibiotics to become widely used. Nowadays, with the increasing incidence of bacterial strains resistant to most classes of antibiotics (5), MBAs have re-emerged as an alternative to prevent and manage infections (6).

Silver, copper and gold are group 11 transition metal elements. These are considered precious and thus have been historically used to mint coins: hence the term *coinage metals*. Their effectiveness (7–12) against different WHO critical priority pathogens (5) has led them to return in popularity as antimicrobials. Silver-based antimicrobials are commercially available and see applications in health including coatings for medical indwelling devices (13), as topical wound dressings (14), and impregnated on textile fabrics (15). Copper is also used in medical settings as part of coatings for high-touch surfaces (10), medical devices (16) and in textiles (17), in addition to a wide variety of usages in agriculture, animal husbandry and aquaculture due to its antifouling, algaecide, antifungal and crop enhancing properties (18). Gold complexes and nanoparticles are under research for their anticancer (19, 20) and antiarthritic (21) properties beyond their bactericidal activity (11, 22, 23), making them good candidates for drug repurposing (24).

Under a One Health perspective, scientists and society must ponder the downstream effects of increased MBA use. The rise of resistance towards metals in the environment is a concern as more MBA applications hit the market, consequence of the acclimation and passive selection of tolerant organisms that find a way to thrive at exposures below lethal concentrations. Instances of co-selection of resistant determinants to both metal stressors and organic antibiotics are now observed (25). Thus, there is a need to look beyond the standard acute toxicity models and investigate bacterial physiology in the context of prolonged exposure to antimicrobials at sublethal levels.

In this context, here we compare *Escherichia coli* bacterial physiology when growing in the presence of sublethal concentrations of silver nitrate (AgNO_3_ – hereby referred to by the elemental symbol Ag), copper sulfate (CuSO_4_ – Cu), and tetrachloroauric acid (HAuCl_4_ – Au). This is achieved by leveraging three different RNA-seq datasets and interrogating the *E. coli* transcriptional regulatory network based on gene expression data. As a result, we present the first transcriptomic comparative analysis of the adaptive and intrinsic response profiles upon acclimation to antimicrobial coinage metal-induced stress. Common and unique gene expression patterns of the response elicited by each metal salt were identified, including increased essential metal uptake (Ag, Cu, Au), cysteine biosynthesis (Cu, Au), change of the metal ion oxidation state (Cu, Au), efflux of metal stressor (Cu), translation and ribosome biogenesis (Au), and cell envelope stress response (Ag).

## RESULTS

### Sublethal Inhibitory Concentration Determination

Susceptibility assays were performed on *E. coli* K12 BW25113. Concentrations of 7, 39 and 10 µM of Ag, Cu and Au, respectively, were deemed as the sublethal inhibitory concentrations to use for the RNA-seq experiments (**Figures S1-S4**). These concentrations exerted a mild inhibitory effect in each bacterial culture as they acclimated to grow in the presence of their respective metal challenges.

**Figure S1.**
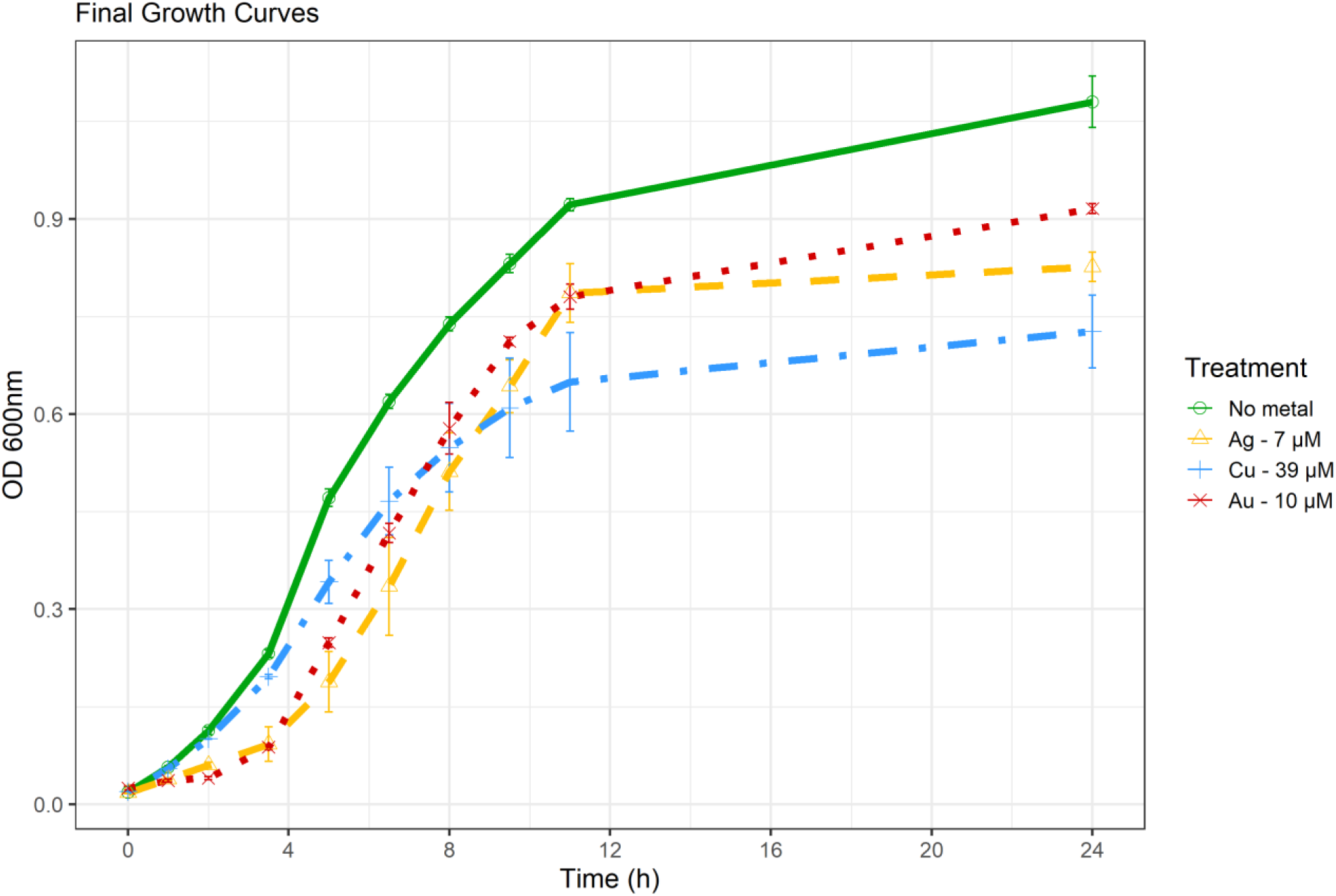
Growth of *E. coli* K12 BW25113 in the presence of sublethal inhibitory concentrations of silver nitrate (Ag), copper sulfate (Cu) and tetrachloroauric acid (Au). Cells were incubated in 250 mL Erlenmeyer flasks at 37 °C 150 rpm using 15 mL of M9-glucose minimal media, spiked with their respective metal salt. Each symbol point is the average of three biological trials. Error bars represent one standard deviation.

**Figures S2-S4.**
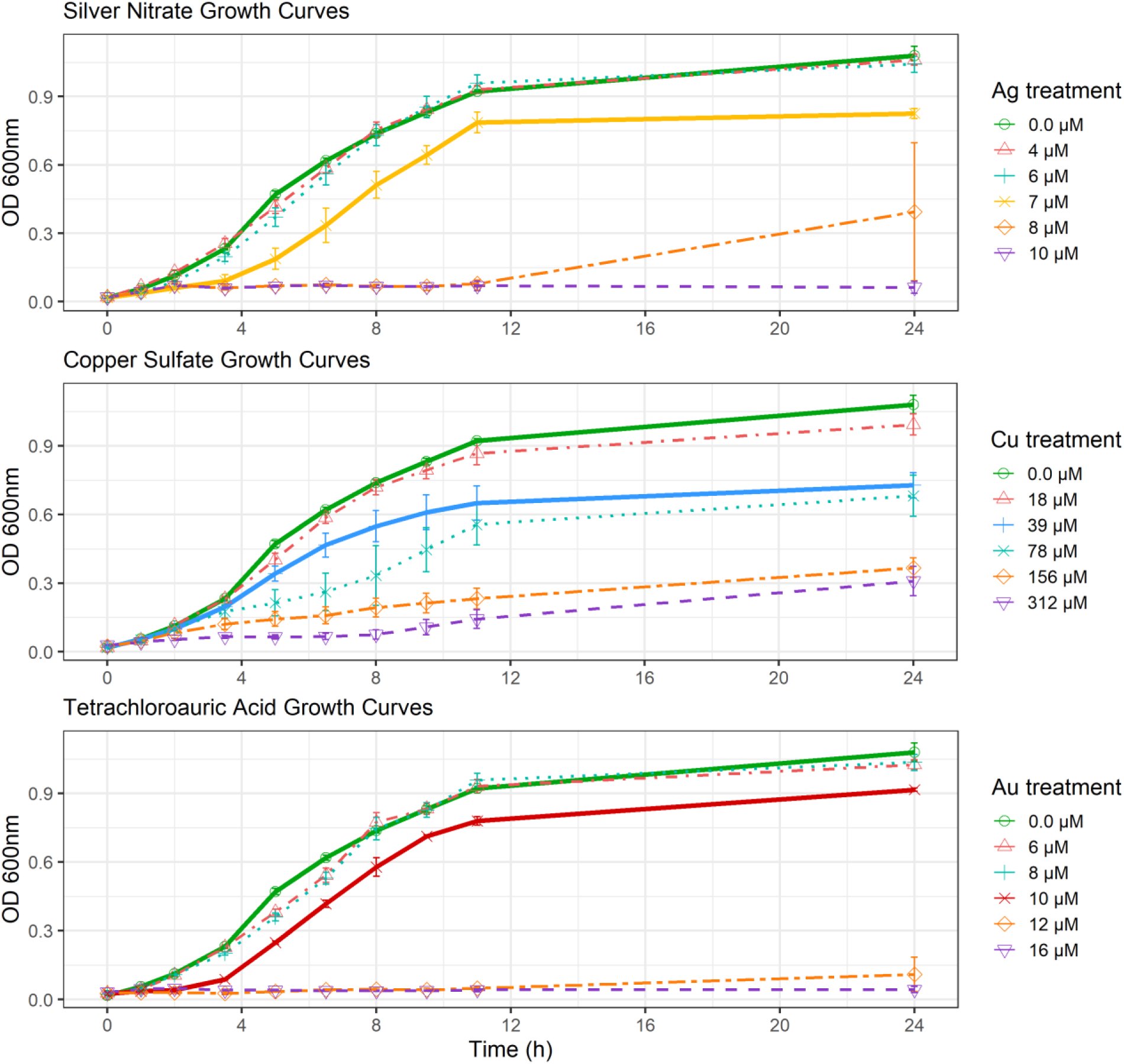
Growth of *E. coli* K12 BW25113 in the presence of coinage metal salts. Cells were incubated in 250 mL Erlenmeyer flasks at 37 °C 150 rpm using 15 mL of M9-glucose minimal media, spiked with their respective metal salt: silver nitrate (Ag, top), copper sulfate (Cu, middle), and tetrachloroauric acid (Au, bottom). Each symbol point is the average of three biological trials. Error bars represent one standard deviation.

### RNA-Seq Data Processing

After processing the sequencing files, differential expression for each metal treatment was obtained by contrasting against their respective non-metal controls. From a total of 4,378 coding sequences in the *E. coli* K12 BW25113 genome assembly, Ag yielded 131 <u>up</u>-regulated <u>d</u>ifferentially <u>e</u>xpressed <u>g</u>enes (UP DEGs) and 558 <u>down</u>-regulated <u>d</u>ifferentially <u>e</u>xpressed <u>g</u>enes (DOWN DEGs); Cu had 80 UP DEGs and 76 DOWN DEGs, while Au did 1028 and 1025 respectively (**Table S1-S3**, **Figures S5-S7**). Principal component analysis was performed on the top 5% most variable genes per dataset after r-log transformation of their expression data, showing clusters of samples according to the treatment used (**Figures S8-S10**).

**Figures S5-S7.**
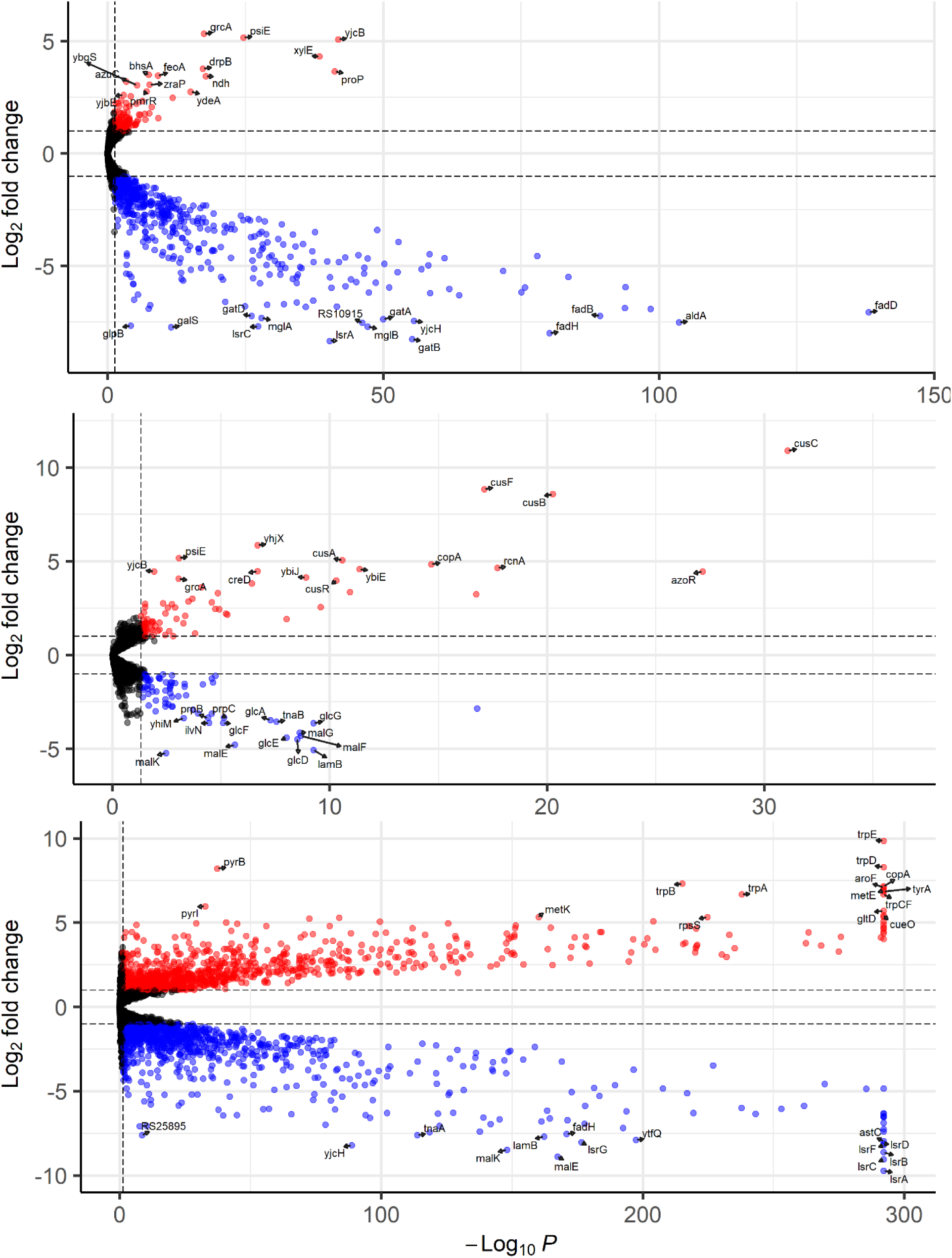
Volcano plots showing differential gene expression of *E. coli* K12 BW25113 after growing in sublethal concentrations of silver nitrate- (top), copper sulfate- (middle) and tetrachloroauric acid- (bottom) spiked M9-glucose minimal media for 10 hours, contrasted against an untreated control. The thresholds for significance were defined by false discovery rate-adjusted p-value < 0.05 and absolute fold-change ≥ 2 (|log_2_ fold change| ≥ 1). Up-regulated differentially expressed genes (DEGs) that exceeded these thresholds are pictured with a red dot, down-regulated DEGs have a blue one; labels with the gene name appear for the top 15 up- and down-regulated DEGs.

**Figure S8.**
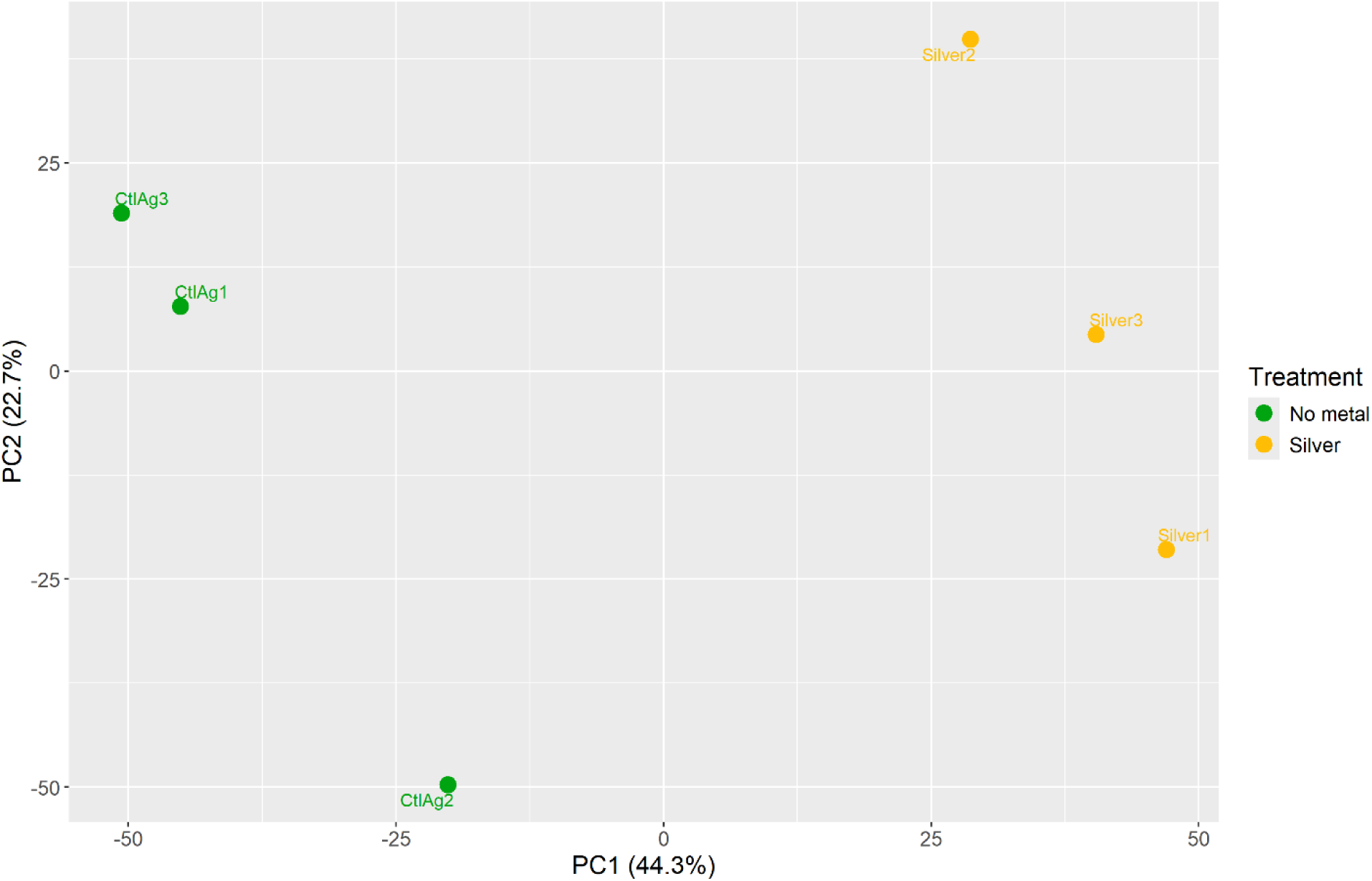
Principal Component Analysis of the samples from the silver nitrate RNA-seq experiment. The r-log transformed raw count values of the 219 most variable genes across samples were used, representing 5% of the 4378 total genes in *E. coli* K12 BW25113. The distances between each point characterize the variation in expression between each sample.

**Figure S9.**
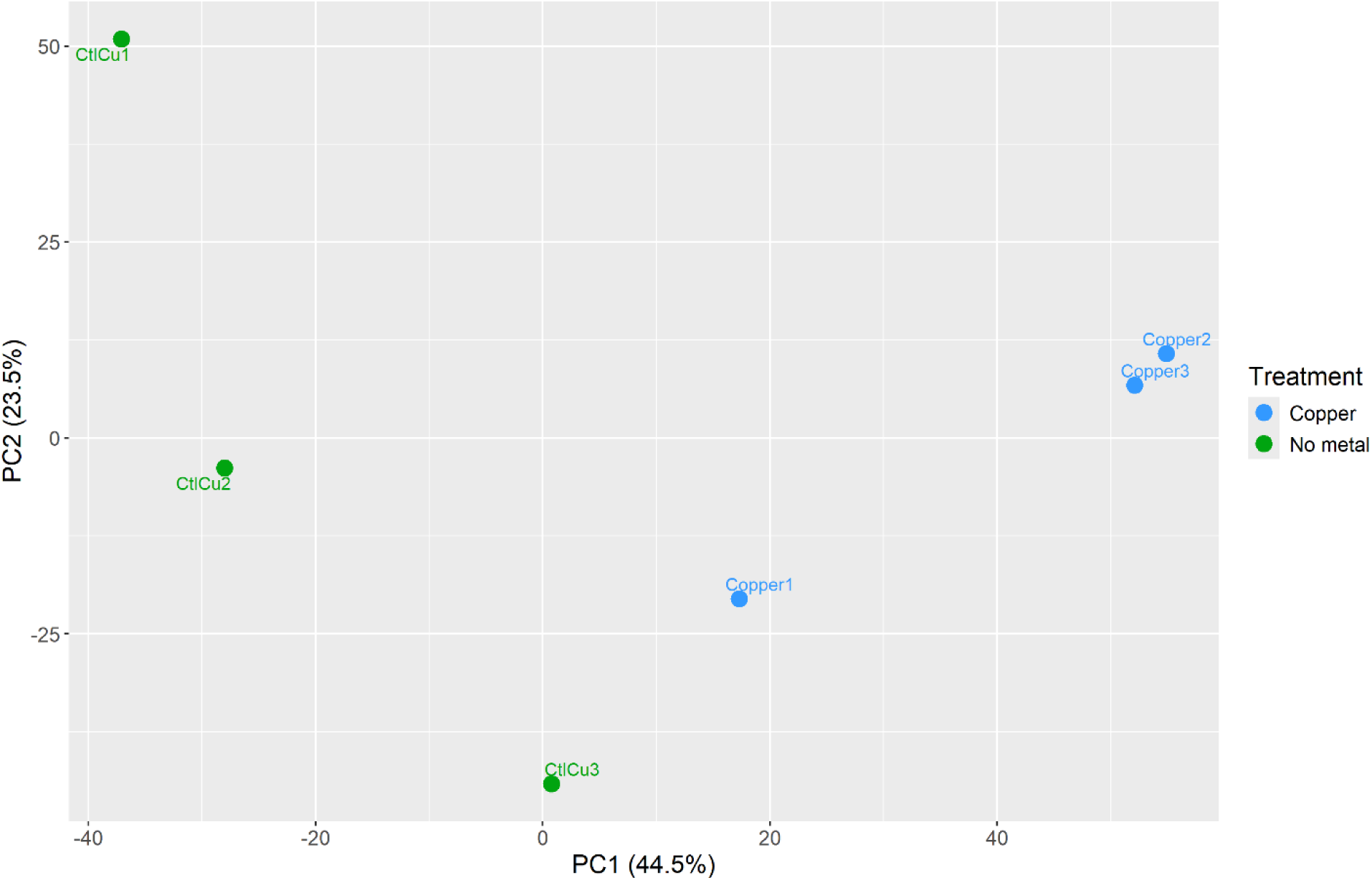
Principal Component Analysis of the samples from the copper sulfate RNA-seq experiment. The r-log transformed raw count values of the 219 most variable genes across samples were used, representing 5% of the 4378 total genes in *E. coli* K12 BW25113. The distances between each point characterize the variation in expression between each sample.

**Figure S10.**
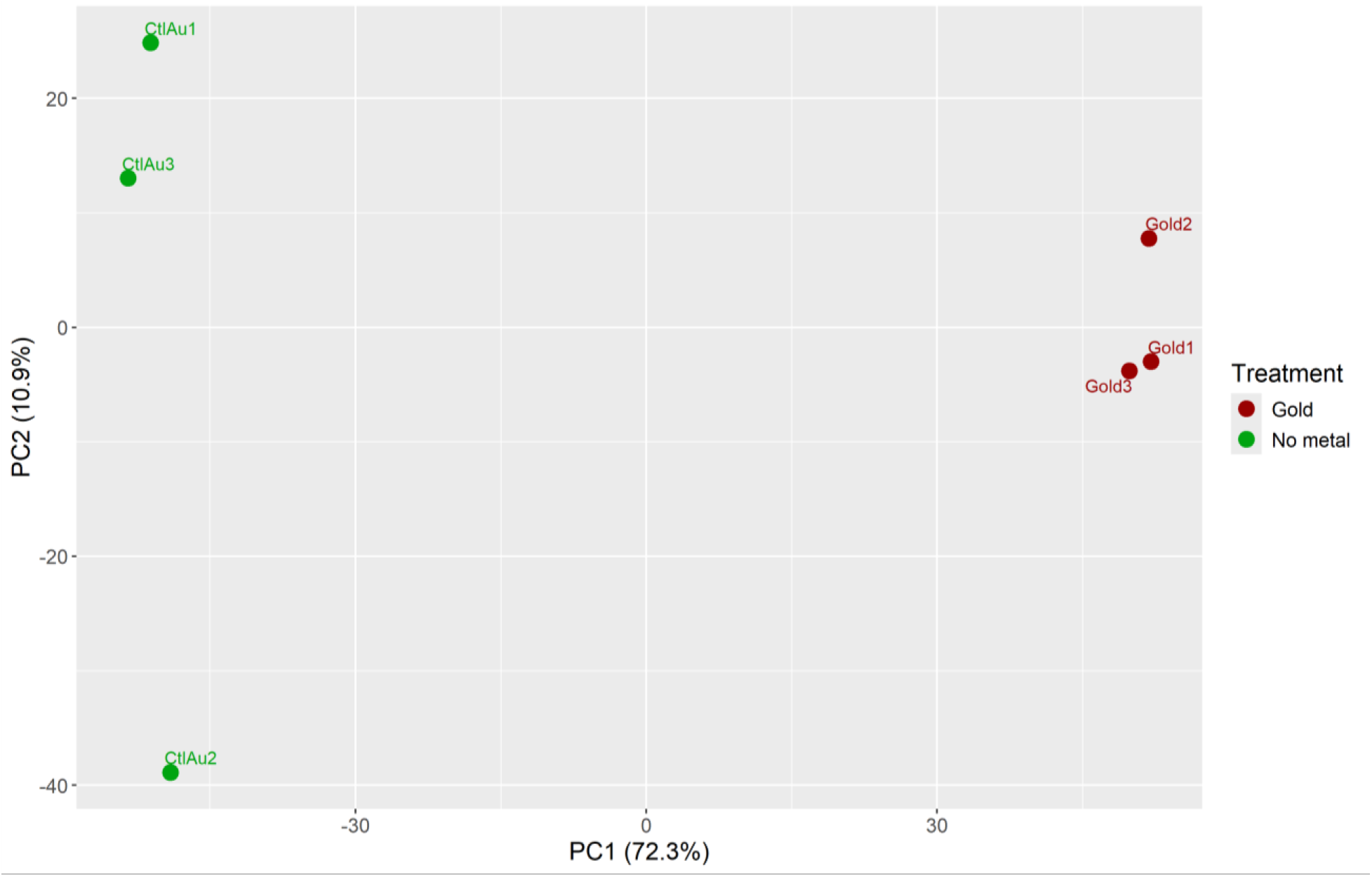
Principal Component Analysis of the samples from the tetrachloroauric acid RNA-seq experiment. The r-log transformed raw count values of the 219 most variable genes across samples were used, representing 5% of the 4378 total genes in *E. coli* K12 BW25113. The distances between each point characterize the variation in expression between each sample.

We generated Venn diagrams to visualize the overlaps in DEGs from our three metal salt datasets (**Figures 1 & 2**). Unique DEGs were found in each of our treatments while also indicating small groups of shared genes between the different metal stressors. Across the three metal salt treatments, 23 genes were shared as UP DEGs (**Table 1**) while 25 were shared DOWN DEGs.

**Figures 1 & 2.**
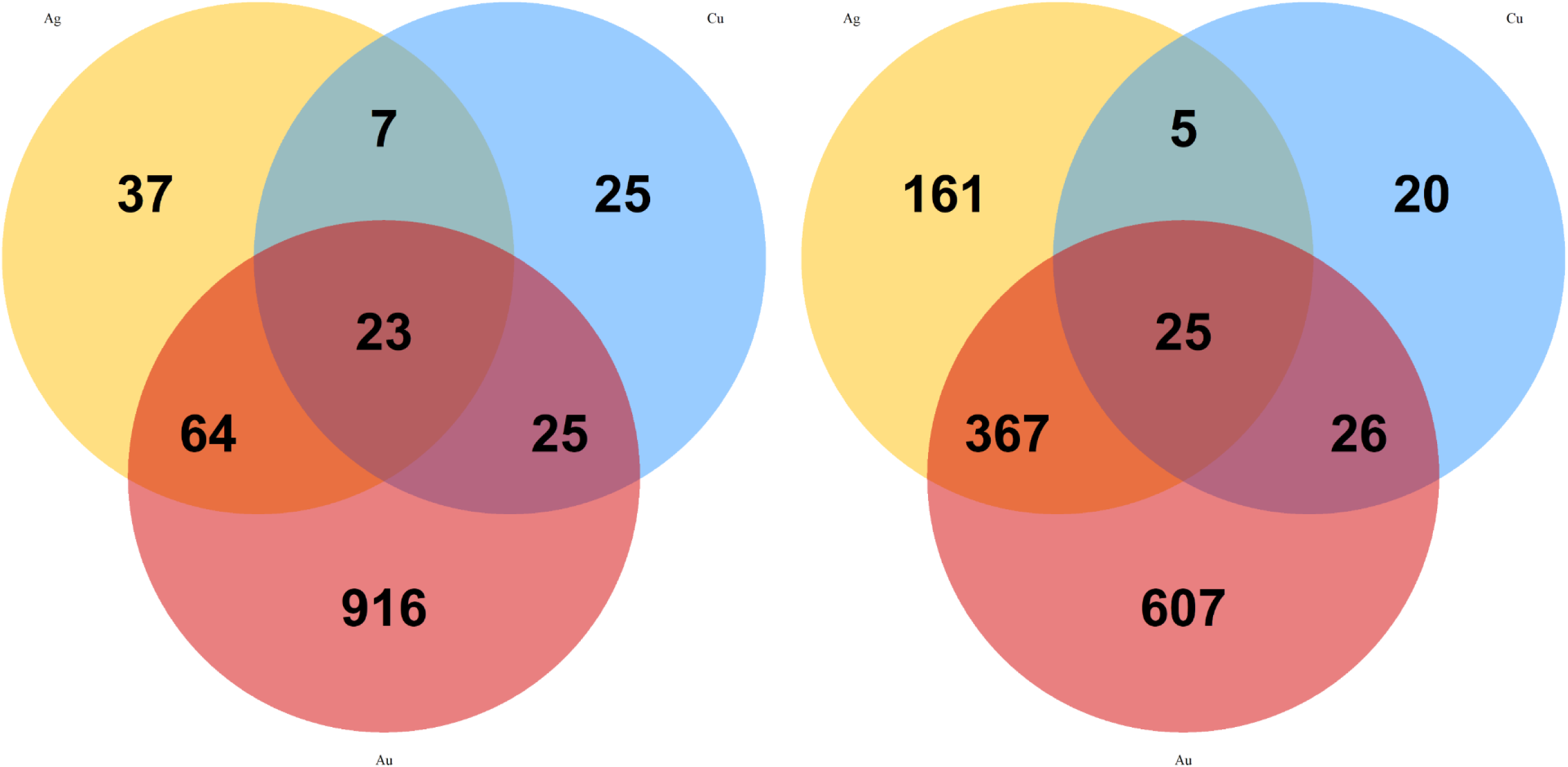
Venn diagrams showing overlaps in up-regulated (left) and down-regulated (right) differentially expressed genes of *E. coli* K12 BW25113 after growing in sublethal concentrations of silver nitrate-, copper sulfate-, or tetrachloroauric acid-spiked M9-glucose minimal media for 10 hours. The yellow circle belongs to the genes from the silver nitrate treatment, blue is for copper sulfate, and red is for tetrachloroauric acid.

**Table 1.** List of the 23 significantly up-regulated genes that overlapped across the silver nitrate, copper sulfate, and tetrachloroauric acid RNA-seq datasets.

| Gene | Product | Uniprot ID |
| --- | --- | --- |
| *ybfA* | YbfA family protein | P0AAU2 |
| *ybiE* | protein YbiE | P0DSE7 |
| *ndh* | NADH-quinone dehydrogenase | P00393 |
| *bhsA* | multiple stress resistance protein BhsA | P0AB40 |
| *azoR* | FMN-dependent NADH-azoreductase | P41407 |
| *ydeA* | L-arabinose MFS transporter | P31122 |
| *ydhR* | monooxygenase | P0ACX3 |
| *tcyP* | L-cystine/L-cysteine shuttle transport system | P77529 |
| *guaB* | IMP dehydrogenase | P0ADG7 |
| *grcA* | autonomous glycyl radical cofactor GrcA | P68066 |
| *serA* | phosphoglycerate dehydrogenase | P0A9T0 |
| *yrbN* | protein YrbN | C1P618 |
| *feoA* | ferrous iron transporter A | P0AEL3 |
| *feoB* | Fe(2+) transporter permease subunit FeoB | P33650 |
| *feoC* | [Fe-S]-dependent transcriptional repressor FeoC | P64638 |
| *glnA* | glutamate--ammonia ligase | P0A9C5 |
| *zraP* | zinc responsive periplasmic chaperone ZraP | P0AAA9 |
| *psiE* | phosphate-starvation-inducible protein PsiE | P0A7C8 |
| *xylE* | D-xylose transporter XylE | P0AGF4 |
| *yjcB* | YjcB family protein | P32700 |
| *proP* | glycine betaine/L-proline transporter ProP | P0C0L7 |
| *fecB* | Fe(3+) dicitrate ABC transporter substrate-binding protein FecB | P15028 |
| *fecA* | Fe(3+) dicitrate transport protein FecA | P13036 |

### Obtaining Biological Significance by Interrogating Regulons of Interest and Co-Expression Patterns

To further investigate the biological context of the gene expression data, we uploaded our raw FASTQ sequencing files to the Integrated System for Motif Activity Response Analysis (ISMARA) (26) web server. With this bioinformatic tool we were able to identify the regulons that had the highest and lowest gene expression activity in our metal salt treatments (**Tables 2-4**), as it detects motifs in the promoters of the *E. coli* K12 genome and infers the activity of gene regulators based on the expression of their target genes. One of the regulons that scored among the top 10 in activity across the three treatments is Fur, whose target genes are known to be involved in maintaining iron homeostasis. Another highlight was the presence of some known copper homeostasis mechanisms, governed by the CusR and CueR regulons, with the latter having high activities in the Cu and Au datasets but not in Ag.

**Table 2.** The top and bottom 10 *E. coli* K12 BW25113 regulators implicated in positive gene expression changes after 10 hours of growth in the presence of sublethal concentrations of silver nitrate, obtained using ISMARA. The z-score is a representation of differential expression induced in the target genes as a number of *n* standard deviations away from zero, with scores indicating up- (z-score > 0) or down- (z-score < 0) regulation in these genes.

| **Regulator** | **Description** | **Z-score** |
| --- | --- | --- |
| Fur | Repressor of ferric uptake regulon. Controls expression of genes involved in iron homeostasis. | **4.748** |
| CpxR | Responds to misfolded periplasmic and inner membrane proteins. | **1.897** |
| Sigma70 | RNA polymerase, sigma D factor. Genes associated with fast cellular growth. | **1.758** |
| PdhR | Activation of fatty acid degradation genes and repression of cell mobility genes. | **1.533** |
| NagC | Coordinates the biosynthesis of D-glucosamine and N-acetylglucosamine with their catabolism. | **1.318** |
| IHF | DNA-bending protein that affects transcription of a wide variety of operons. | **1.123** |
| MetJ | Represses the methionine operon. | **1.082** |
| PhoP | Regulates genes involved in magnesium uptake, acid resistance and lipopolysaccharide modification. | **1.054** |
| GadE | Central activator of the acid response system. | **0.916** |
| Cra | Helps modulate the flow of central carbon metabolism. | **0.854** |
| GalR | Repressor of transport and catabolism of D-galactose. | **-1.197** |
| MalT | Activator of the maltose regulon: uptake and catabolism of malto-oligosaccharides. | **-1.214** |
| UxuR | Repressor of transport and catabolism of β-D-glucuronides, glucuronate, and gluconate. | **-1.227** |
| GlpR | Repressor of transport and catabolism of glycerol-3-phosphate. | **-1.302** |
| Sigma54 | RNA polymerase, sigma N factor. Genes associated with nitrogen-related processes. | **-1.340** |
| TorR | Activates the trimethylamine-N-oxide anaerobic respiratory system. | **-1.359** |
| FlhDC | Regulates flagella formation, operation, and carbon source metabolism. | **-1.371** |
| FadR | Regulates fatty acid degradation while it activates fatty acid biosynthesis. | **-3.648** |
| ArcA | Repressor of a wide variety of aerobic enzymes under anaerobic conditions. | **-4.982** |
| CRP | Global activator of secondary carbon source pathways. | **-13.487** |

**Table 3.** The top and bottom 10 *E. coli* K12 BW25113 regulators implicated in positive gene expression changes after 10 hours of growth in the presence of sublethal concentrations of copper sulfate, obtained using ISMARA. The z-score is a representation of differential expression induced in the target genes as a number of *n* standard deviations away from zero, with scores indicating up- (z-score > 0) or down- (z-score < 0) regulation in these genes.

| Regulator | Description | Z-score |
| --- | --- | --- |
| CusR | Copper extrusion system CusCFBA. | 1.986 |
| YedW | Hydrogen peroxide response regulator. | 1.986 |
| CueR | Controls the expression of the *copA* and *cueO* copper homeostasis genes. | 1.851 |
| CysB | Activator of sulfur metabolism and cysteine biosynthesis. | 1.584 |
| PhoB | Activates expression of the *pho* regulon in response to environmental inorganic phosphate. | 1.398 |
| Sigma70 | RNA polymerase, sigma D factor. Genes associated with fast cellular growth. | 1.308 |
| FNR | Mediates transition from aerobic to anaerobic growth. | 1.226 |
| PurR | Regulates de novo synthesis of purine and pyrimidine nucleotides. | 1.205 |
| RcnR | Represses the expression of the cobalt and nickel metallo-regulator operon RcnAB. | 1.117 |
| Fur | Repressor of ferric uptake regulon. Controls expression of genes involved in iron homeostasis. | 0.962 |
| GlcC | Repressor of transport and catabolism of glycolate. | -0.929 |
| Fis | DNA-bending protein that affects transcription of a wide variety of operons. | -1.010 |
| Nac | Regulates genes involved in nitrogen metabolism under nitrogen-limiting conditions. | -1.070 |
| GadW | Regulates expression of acid resistance genes *gadA* and *gadBC*. | -1.202 |
| GadX | Activates genes involved in acid resistance, such as *gadA* and *gadBC*. | -1.216 |
| FadR | Regulates fatty acid degradation while it activates fatty acid biosynthesis. | -1.285 |
| LsrR | Represses transcription of the *lsr* operon, belongs to the quorum-sensing system. | -1.489 |
| Sigma38 | RNA polymerase, sigma S factor. Genes associated with stationary phase. | -1.690 |
| TorR | Activates the trimethylamine-N-oxide anaerobic respiratory system. | -1.749 |
| MalT | Activator of the maltose regulon: uptake and catabolism of malto-oligosaccharides. | -2.558 |

**Table 4.**
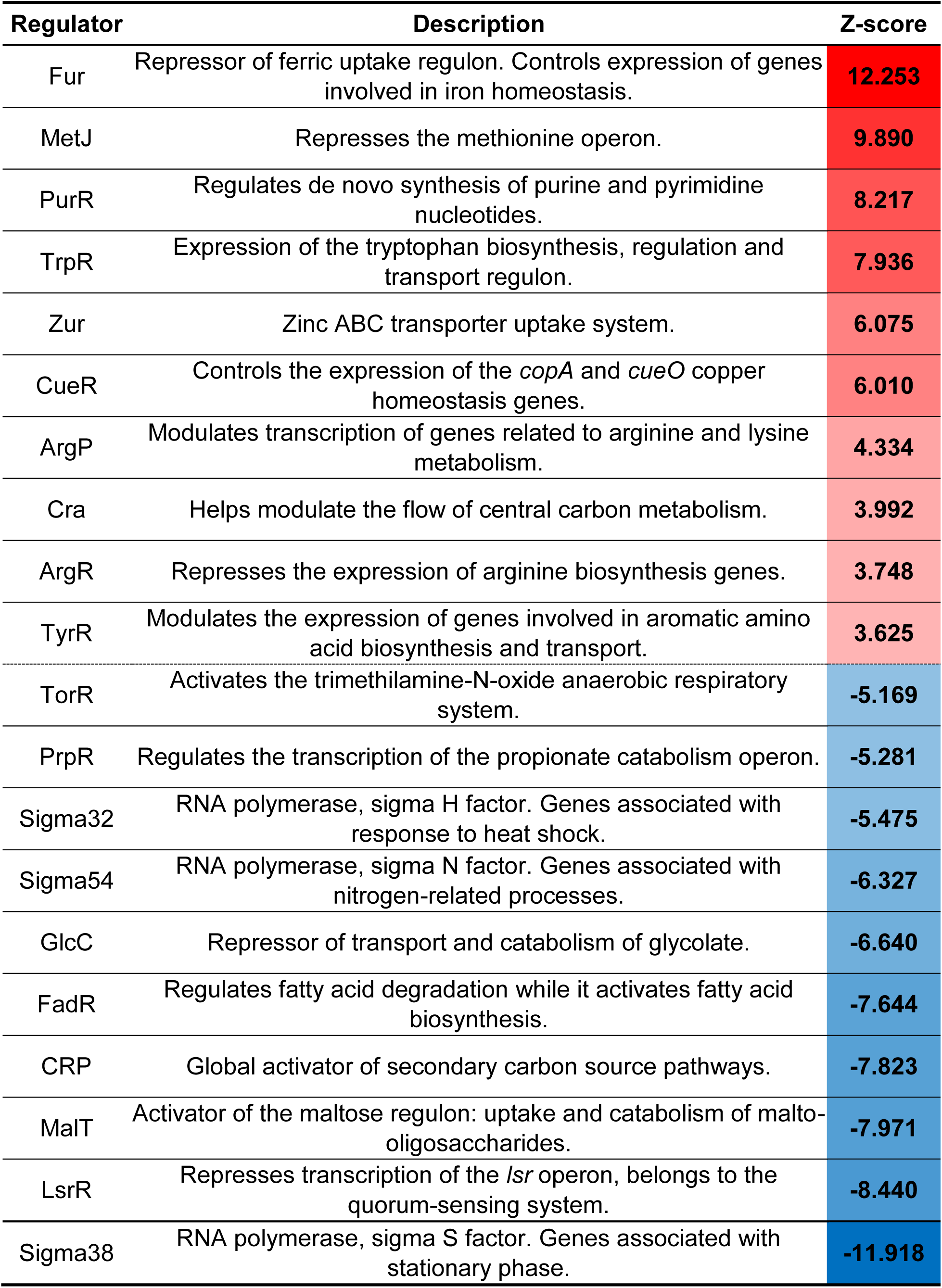
The top and bottom 10 *E. coli* K12 BW25113 regulators implicated in positive gene expression changes after 10 hours of growth in the presence of sublethal concentrations of tetrachloroauric acid, obtained using ISMARA. The z-score is a representation of differential expression induced in the target genes as a number of *n* standard deviations away from zero, with scores indicating up- (z-score > 0) or down- (z-score < 0) regulation in these genes.

To identify co-expression patterns, we queried an *E. coli* transcriptional regulatory network based on independently modulated signals, known as iModulons (27). For this purpose, we utilized the *E. coli* PRECISE-1K dataset (28). This is composed of 201 groups of independently <u>modul</u>ated groups of genes identified by an unsupervised machine-learning algorithm that have shown clear co-expression patterns across 533 different experimental conditions. Each iModulon is subsequently linked to the effect of a single, multiple, or no known regulators. After determining the expression activities for each of the 201 iModulons in our experimental conditions (**Figures 3-6**), we discovered that the Au treatment yielded the highest amount of significant iModulons, with 38; meanwhile, Ag and Cu reported 13 and 8 significant iModulons, respectively. There were no shared significantly up-regulated iModulons across the three metal treatments, which is an indicator of how each metal stressor required a different type of response for the cultures to reach acclimation.

**Figures 3-6.**
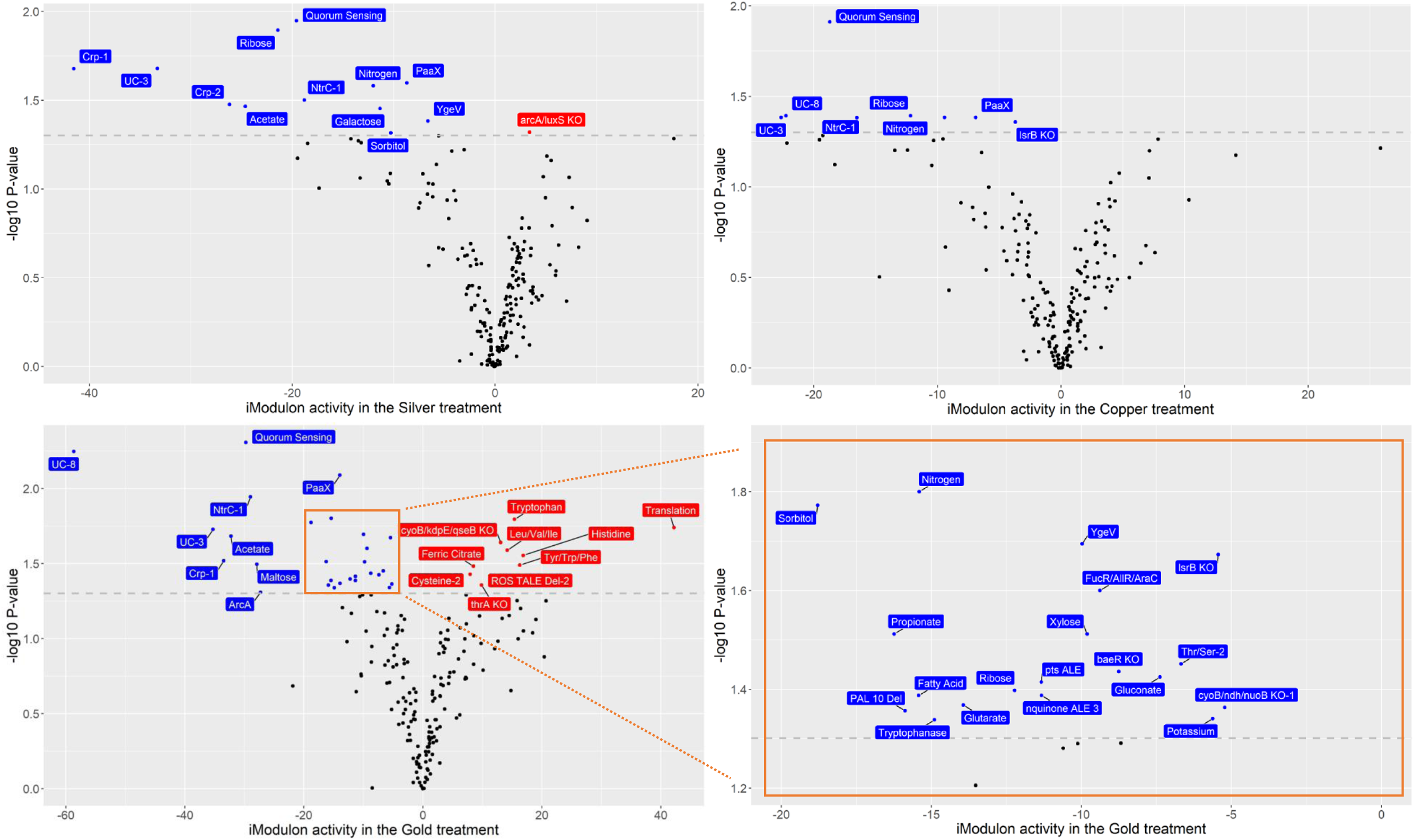
Distribution of iModulon activities in the silver nitrate (top left), copper sulfate (top right) and tetrachloroauric acid (bottom left and right) datasets relative to their non-metal controls. Positive activities in the x-axis indicate iModulons that are more active in the metal condition compared to the control, and vice versa. Statistical significance was calculated using the cumulative log-normal distribution of each iModulon across all conditions. The highlighted data points correspond to iModulons with a p-value score lower than 0.05.

## DISCUSSION

Bacterial susceptibility studies have traditionally focused on acute toxicity models, where the focus is to measure lethality and response over short periods of time (0 to 4 hours) after exposing a growing culture to a bactericidal concentration of an antimicrobial. This has the downside effect of ignoring chronic exposure models where bacterial physiology is modified as the organism acclimates to the antimicrobial stress. Nowadays, as the scientific community becomes aware of the One Health perspective (29), these physiological adaptations that lead to antimicrobial tolerance in prolonged exposure conditions have gained interest. Here, we analyze the adaptive gene expression of *E. coli* growing under sublethal inhibitory antimicrobial coinage metal stress. By leveraging regulon expression analysis and machine learning-based clusters of co-expressed genes, we compared shared and unique trends in the bacterial physiology acclimated to sustained Ag, Cu, and Au-induced stress. Our findings are summarized in **Figure 7** and discussed below.

**Figure 7.**
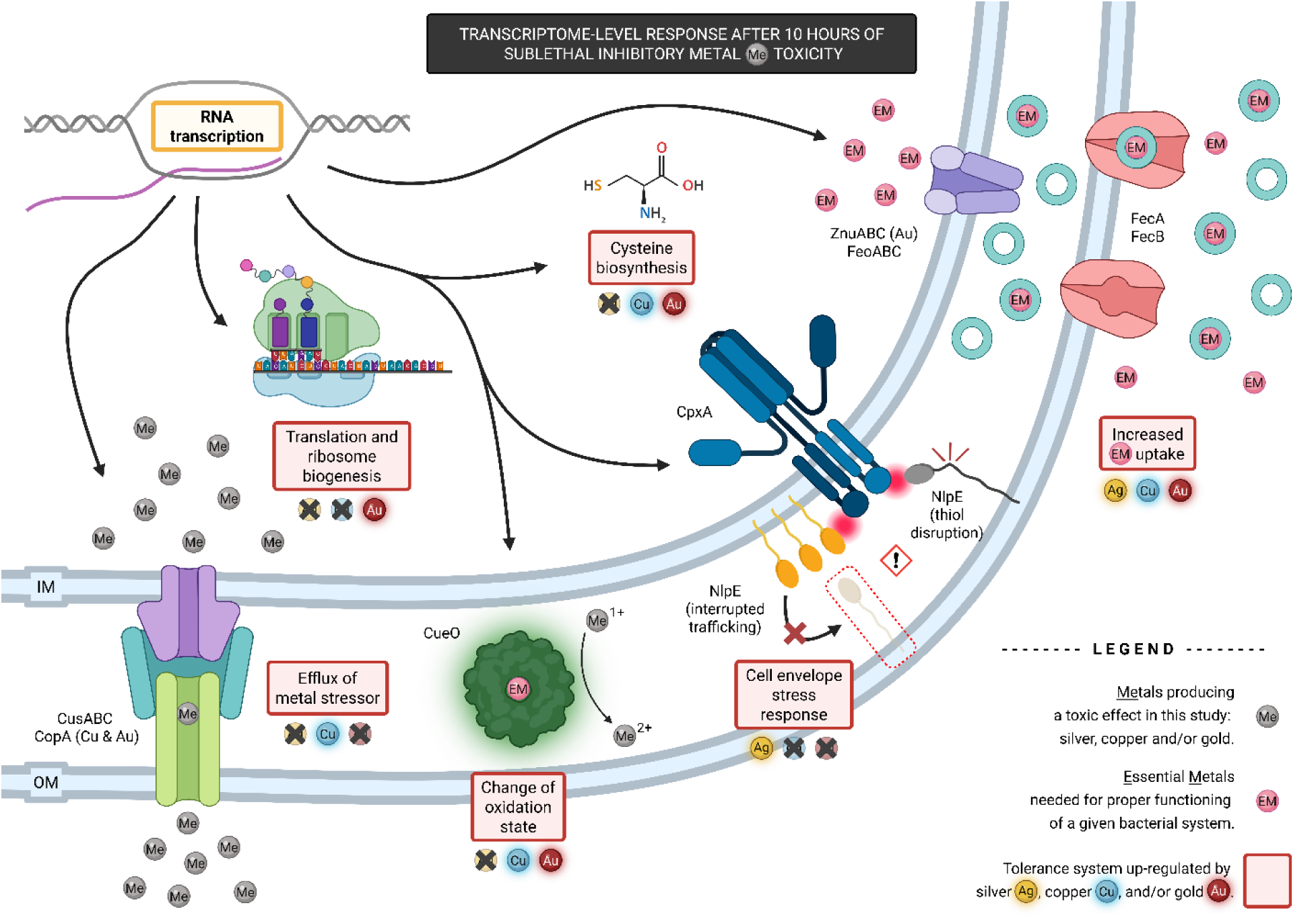
Generalized model of the *E. coli* K12 BW25113 transcriptome-level response to sublethal inhibitory stress induced by antimicrobial coinage metal salts after 10 hours of growth. An increase in the expression of essential metal (EM) uptake systems was observed: iron metabolism and uptake were affected by each metal, while the ZnuABC zinc uptake system was up-regulated by gold (Au). Silver (Ag) triggered the cell envelope stress response which acts upon outer membrane protein trafficking disruption and misfolding (through the NlpE sentry protein (30)). Copper (Cu) activated the expression of the Cus system, which effluxes excess copper ions. The other copper homeostasis system in *E. coli*, Cue, is sensitive to monovalent metal cations (31) and was up-regulated by Au and Cu: CueO specializes in changing the oxidation state of the more toxic Cu(I) to the less toxic Cu(II), while CopA exports Cu(I) to the periplasm. Cysteine biosynthesis and sulfur metabolism were induced by Cu and Au to replenish the antioxidant pool of the cell and maintain redox balance in the cytoplasm and periplasm. Au also caused increased gene expression activity in ribosome biogenesis and protein translation-related genes. Created with BioRender.com.

From a chemistry perspective, we see both similarities and differences of these coinage metal elements. As one goes down group 11 in the periodic table (copper ◊ silver ◊ gold), ionization energy and atomic radius increase (giving higher coordination number), while redox chemistry tends to decline to the point that gold is often considered chemically inert (32). Typically, the enthalpies of atomization of the elements increase the lower you go in each group, but differences tend to be small in the enthalpies between the 3d, 4d, and 5d orbitals of these group 11 elements, reflecting similar strengths of bonding. However, the *ns*^1^ outer electron of these coinage metals can be ionized at higher energies over the alkali metals. Copper and silver have similar ionization energies at ∼8 eV but gold is higher at ∼9.5 eV. These parameters would reflect the on-off bonding rates with the various soft biochemical bases, particularly thiol groups. Regardless, we would expect the cupride, argentide and auride ion forms to be more reactive than their cuprate, argenate and aurate forms of copper, silver, and gold, respectively. Thus, their toxicity may depend more on the redox potential and pH of the cell (in relation to the Pourbaix diagram for each element), which would dictate the relative concentration of the -ide ion forms in the cell.

Another consideration in coinage metal chemistry is the flexibility of their ligand coordination geometry. Concerning copper, the Irving-Williams series have it as the most promiscuous regarding its ligands and geometry, which allows it to effectively compete for metal ligand sites in proteins and other biomolecules, whereas gold is more ridged and less likely to outcompete essential metal sites (33). This would give the theoretical order of toxicity of the coinage metal elements as follows: copper > silver > gold. Yet, silver has the highest level of toxicity because cells have evolved important dedicated homeostasis systems to protect cellular targets from copper, which do not exist for the other two.

### Coinage metal salts affect the expression of iron homeostasis genes

Based on the 23 UP DEGs that were shared between the three metal salt treatments, we interrogated the Gene Ontology (GO) enrichments of this subset using the Search Tool for the Retrieval of Interacting Genes/Proteins (STRING) (34). This resulted in three Biological Process GO terms being enriched: transition metal ion homeostasis (GO:0055076), iron import into cell (GO:0033212), and iron ion homeostasis (GO:0055072) (**Figure 8**). As such, it is natural to infer a disruption of iron homeostasis and metabolism is being caused by the three metal stressors in question. This is further supported by the overall increase in gene expression activity within the Fur regulon (ferric <u>u</u>ptake regulation, **Figure S11**), which features as one of the regulons with the highest activity across all metal treatments (#1 in Ag and Au, #10 in Cu) (**Tables 2-4**).

**Figure 8.**
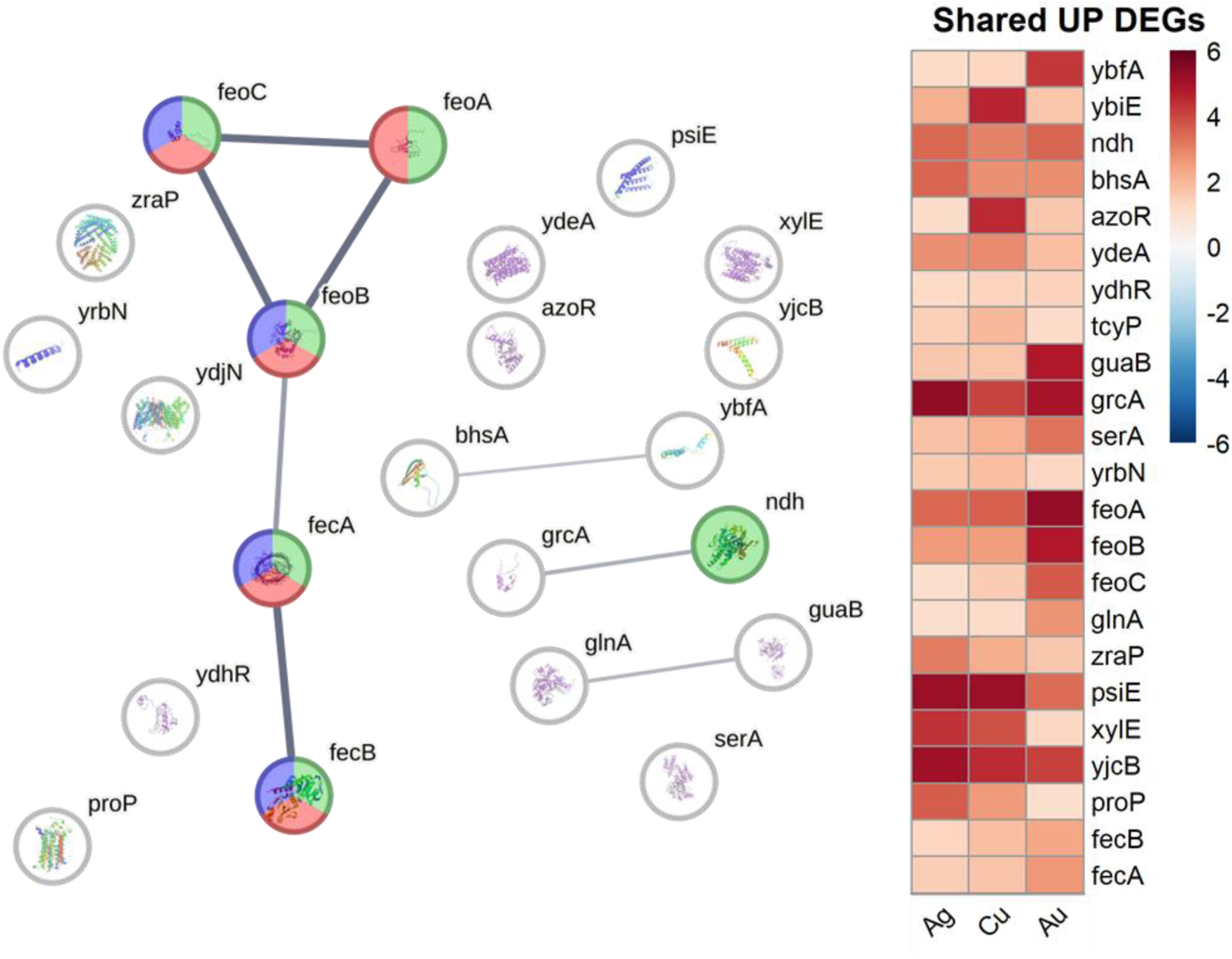
Network interactions map (left) and average log_2_-fold change in gene expression for the 23 shared significant up-regulated genes between the silver nitrate (Ag), copper sulfate (Cu), and tetrachloroauric acid (Au) conditions (right). Each node from the network map represents a gene, and these are connected by edges to adjacent nodes representing a possible interaction between them. A thicker edge between nodes indicates a higher degree of confidence of that interaction. Certain nodes are colored based on the enriched Gene Ontology term they are a part of: green for transition metal ion homeostasis (GO:0055076), blue for iron import into cell (GO:0033212), and red for iron ion homeostasis (GO:0055072). Each metal salt treatment was contrasted against a non-metal challenge control, with three biological trials each.

**Figure S11.**
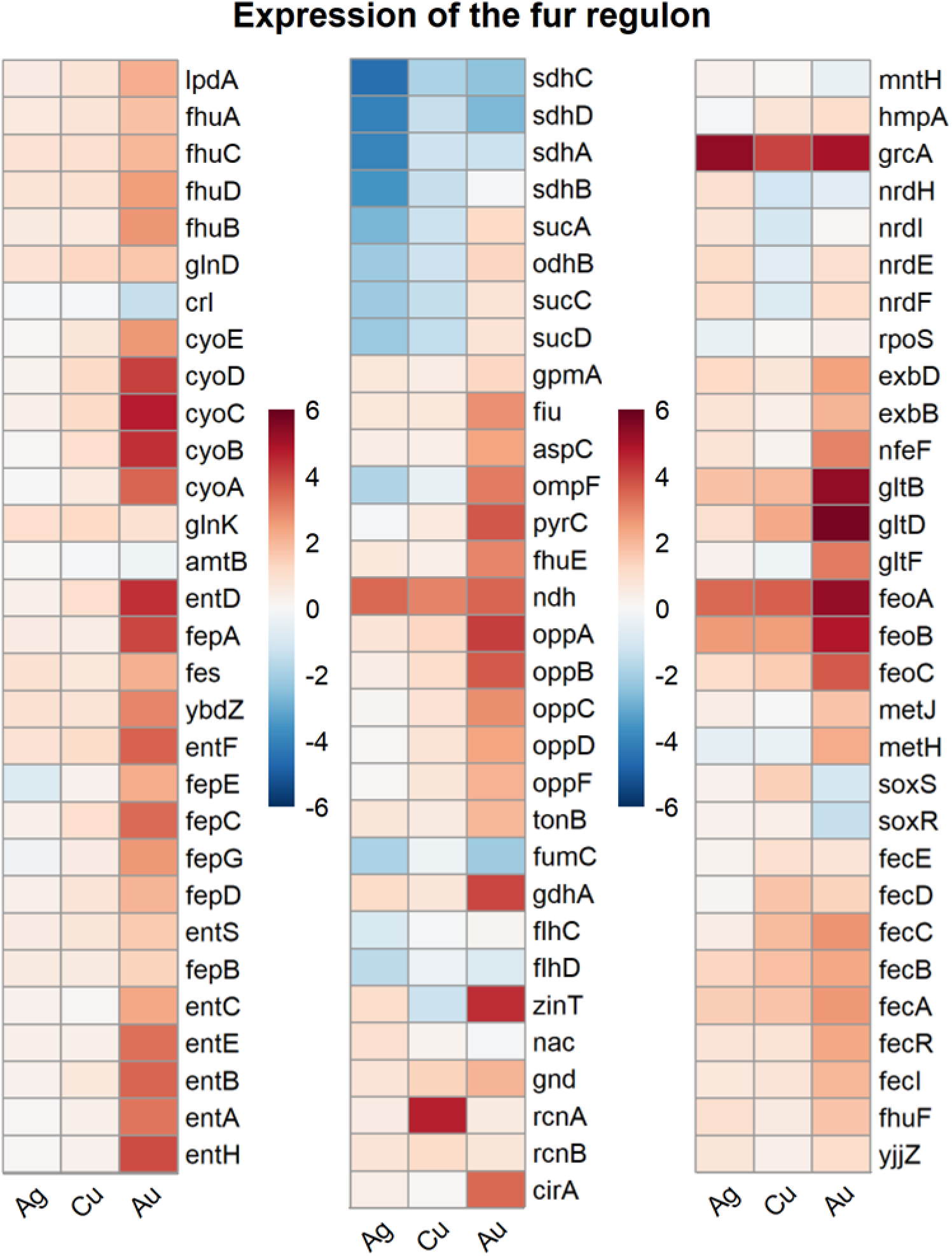
Average log_2_-fold change in gene expression for genes that are part of the Fur regulon. Each column corresponds to a different experimental condition: silver nitrate (Ag), copper sulfate (Cu), and tetrachloroauric acid (Au). Every experiment was contrasted against a non-metal challenge control, with three biological trials each.

Fur is a transcriptional regulator that can bind a 2-iron, 2-sulfur cluster ([2Fe-2S]). It has been proposed that its main regulatory role is to act as a sensor of ferrous iron (Fe(II)) within the cell (35). When there is an abundance of intracellular Fe(II), [2Fe-2S] centres become more available and trigger a conformational change in Fur (36), enabling its genetic repressor role. Conversely, when Fe(II) is scarce, Fur repression is alleviated, resulting in a higher expression of its target genes. Seven Fur target genes over four transcriptional units stand out as UP DEGs across the three conditions: NADH-quinone dehydrogenase *ndh*, autonomous glycyl radical cofactor *grcA*, ferrous iron transport system *feoABC*, and ferric (Fe(III)) citrate transport components *fecAB*. The presence of these last two iron import operons would indicate that the three metal salts are inducing an iron-deprived state in their respective cultures. This type of genetic response has also been observed with gallium nitrate (37).

While it is a shared trend, it is possible that each coinage metal elicited this type of response through different mechanisms. Ag(I) is known to attack exposed [Fe-S] clusters (38) given its nature as a soft acid, interacting with thiol ligands (39). Copper, being able to switch between its soft acid cuprous (Cu(I)) and borderline acid cupric (Cu(II)) forms under cellular redox potentials (40), has the ability to compete iron out from these clusters. This results in an increase in reactive oxygen species (ROS) as the free iron undergoes Fenton chemistry, creating a cycle of dysruption as ROS can target other [Fe-S] centers in the cell. On the other hand, tetrachloroauric acid presents gold in its trivalent oxidation state. Being a harder acid, Au(III) wouldn’t exactly fit Ag(I) or Cu(II)’s models, but it’s been reported that it indirectly triggers a major oxidative imbalance in *E. coli*, characterized by an increase in superoxide (11). This may be the result of Au(III) intracellular reduction to Au(I), which would allow it to also compete with Cu(I) sites and possibly trigger Fenton chemistry. Such a phenomenon could be explained by a reduction of Au(III) to Au(I) caused by an undetermined cellular redox activity, which would allow it to compete with Cu(I) sites and possibly trigger Fenton chemistry. It’s been suggested that a local production of Au(I) salts could happen in a reductive environment such as the *E. coli* cell (41), and metal reduction towards their elemental state is one of the bacterial responses to metal stress (42). This could partly explain the indirect disruption of iron homeostasis and metabolism by Au, although the exact mechanism is yet to be elucidated.

### Expression of the Cus and Cue systems is heavily influenced by the presence of copper

The ability of copper to act as a soft or borderline acid by cycling between Cu(I) and Cu(II) grants a wide range of biochemical applications for this metal. The former tends to have more affinity for soft bases (thioethers and thiols), while the latter shows increased affinity for borderline bases (imidazole nitrogen groups, glutamate, aspartate) (43). As such, it acts as a cofactor in metalloproteins with various functionalities, including oxidation-reduction reactions, electron transfer and transport, and denitrification (44). This feature brings forward a dilemma – copper is an essential trace element, but too much copper becomes toxic due to a) its tendency to displace other essential metals from their ligands (such as zinc and iron) (45, 46), and b) its ability to catalyze hydroxyl radical formation through Fenton chemistry (47, 48).

To protect against copper toxicity, microorganisms have evolved mechanisms that directly control its homeostasis. *E. coli* relies on two main systems to achieve this, each regulated separately. The cytoplasmic CueR requires two Cu(I) ions to become active and promote the transcription of *cueO* and *copA* (49), acting as a de-facto Cu(I) sensor in the cytoplasm. This triggers a detoxifying event in which the P-type ATPase CopA and its Cu(I) cytoplasmic chaperone CopA(Z) isoform works in tandem to efflux Cu(I) ions to the periplasm (50–53), where they are oxidized to the less toxic Cu(II) form by CueO (54). On the other hand, the two-component signal transduction system CusSR senses increases in the periplasmic concentration of Cu(I) and promotes the transcription of the Cus system (49, 55), comprised of the *cusCFBA* and *cusSR* transcriptional units. The RND efflux complex CusABC extrudes Cu(I) ions accumulated in the periplasm and cytoplasm (56). It is assisted by the Cu(I) chaperone CusF, which works in tandem with CopA to transfer heavy metal ions to CusABC (57).

In our experiment, Cu stress caused a notable increase in gene expression for both copper homeostasis genes in the CueR regulon, as expected (**Table 3**). Surprisingly, Au presented an even sharper increase, while Ag only increased *copA* expression marginally (**Figure 9**). Meanwhile, the CusR regulon was significantly up-regulated by Cu, with only CusC being an UP DEG in the Ag treatment.

**Figure 9.**
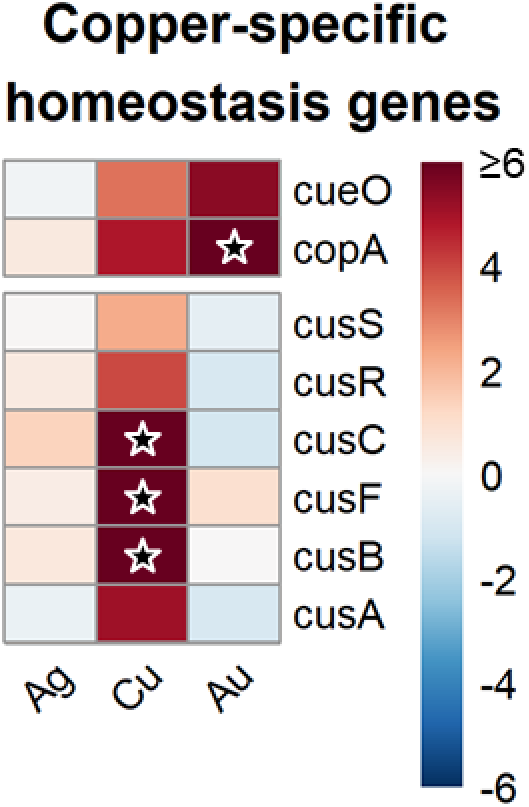
Average log_2_-fold change in gene expression for genes that are known to play a central role in maintaining copper homeostasis in *E. coli* K12 BW25113. The first pair of genes are regulated by CueR, while the second cluster is regulated by CusR. Each column corresponds to a different experimental condition: silver nitrate (Ag), copper sulfate (Cu), and tetrachloroauric acid (Au). Every experiment was contrasted against a non-metal challenge control, with three biological trials each. A ★ represents a log_2_-fold fold change value greater than 6.

There is evidence in the literature of CueR responding to the presence of Ag, Cu, and Au metal salts (31, 41, 58), even if the copper-detoxifying functions of CueO or CopA are not necessarily compatible with silver (59) or gold (58), respectively. This is notable because CueR is reported to have a high selectivity for monovalent metal cations (31), which would fit with the possibility of a localized production of Au(I) salts within the reductive environment of an *E. coli* cell (41). On the other hand, the increase in gene expression of the CusR regulon being caused by Cu is a validation of copper efflux being one of the essential mechanisms to cope with an excess of this metal. While it’s been reported that the Cus system can also respond to Ag(I) ions (60) and efflux them (61), the increase in expression activity by the Ag treatment was markedly smaller than that of Cu exposure. Beyond copper homeostasis, our results add nuance to the role of the Cue and Cus systems when facing Ag and Au stress.

### Silver activates the Cpx cell envelope stress response

One of the unique signals we detected in our Ag experiment was the increased gene expression activity in the CpxR regulon. The Cpx response is composed of a two-component signal transduction system (the sensor kinase CpxA and the transcriptional regulator CpxR), and senses misfolding and misplacement of proteins in the cell envelope (62). Multiple stimuli have been documented to induce it, including acute exposure to copper chloride (CuCl_2_) (63, 64). A model has been proposed where Cu(I) inhibits lipoprotein maturation while they are in traffic from the inner membrane (IM) to the outer membrane (OM). This prevents the sentry lipoprotein NlpE from reaching the OM, causing it to accumulate in the IM which ultimately triggers the Cpx response (30).

While Ag toxicity is known to attack the cell envelope (65), we believe this is the first report of an up-regulation of the CpxR regulon by this metal stressor. Eight CpxR-regulated genes were above significance thresholds in our Ag experiment: periplasmic serine endoprotease *degP*, hemolysin expression modulator *hha*, 3-deoxy-7-phosphoheptulonate synthase *aroG*, curli assembly components *csgFE*, multiple antibiotic resistance transcriptional regulator *marR*, predicted inner membrane protein *yqaE*, and Cpx response modulator *cpxP*. All of these demonstrated slight increases in expression in the Cu dataset, with *csgFE* and FtsH protease modulator *yccA* being UP DEGs as well (**Figure 10**). Considering that both Cu(I) and Ag(I) ions (soft acids) will tend to attack thiol groups from cysteine residues (soft base) (39), it is theoretically possible for both to oxidize thiols to induce this type of response. Nonetheless, this induction of the Cpx response from Ag supports cell envelope homeostasis to be a target for silver ion toxicity.

**Figure 10.**
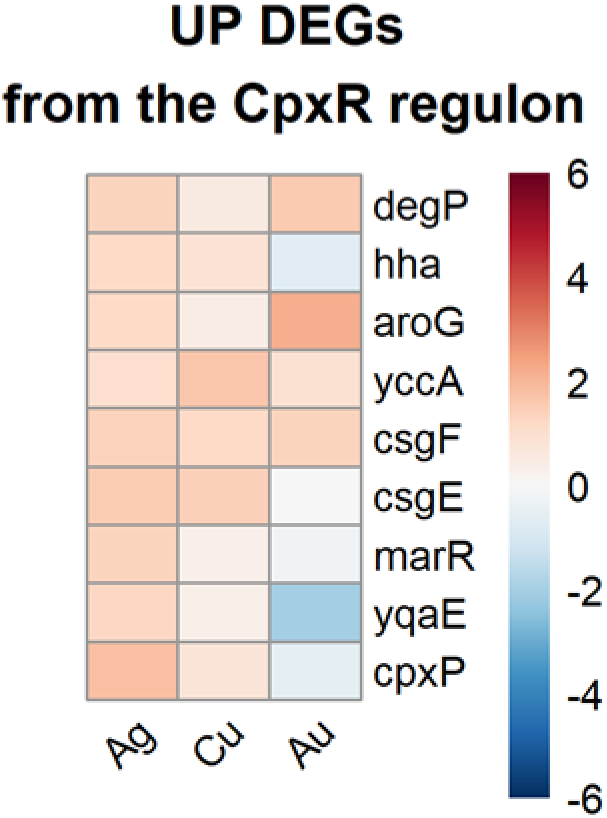
Average log_2_-fold change in gene expression for CpxR-regulated differentially expressed genes for the Ag and Cu treatments in *E. coli*. Each column corresponds to a different experimental condition: silver nitrate (Ag), copper sulfate (Cu), and tetrachloroauric acid (Au). Every experiment was contrasted against a non-metal challenge control, with three biological trials each.

### Gold affects the expression of genes related to sulfur metabolism, translation, ribosome biogenesis and zinc uptake

Previous reports have identified oxidative stress as one of Au’s sources of toxicity, causing redox imbalance and a depletion of reduced thiols (11, 66). This may be reflected in metal–sulphur bond stability: the Cu-S bond is slightly less stable than Au-S, while the Ag-S bond is very stable. For Au-S, this would imply that rather than a reaction stopping at a M + RSH <-> M-S-R, it continues to 2MSR -> RS-SR + 2M. For this reason, we decided to look at the transcription activities in the CysB regulon. This transcription factor regulates genes involved in pathways such as cysteine biosynthesis and sulfate assimilation, having a central role in replenishing the pools of antioxidant molecules such as glutathione and thioredoxin (67). There is an overall trend of up-regulation across CysB’s target genes in the Au treatment (**Figure 11**), on top of the significant up-regulation of *trxB* (log_2_-fold change: 2.25), which codes for thioredoxin reductase and is a known target for auranofin toxicity (66), and the *gsiABCD* transcriptional unit (average log_2_-fold change: 1.44) that codes for a glutathione import system. The latter is not part of the CysB regulon but is often co-expressed with some of those genes as part of the Cysteine-1 iModulon (27). This type of response is consistent with the premise of Au affecting redox balance and reduced thiols balance.

**Figure 11.**
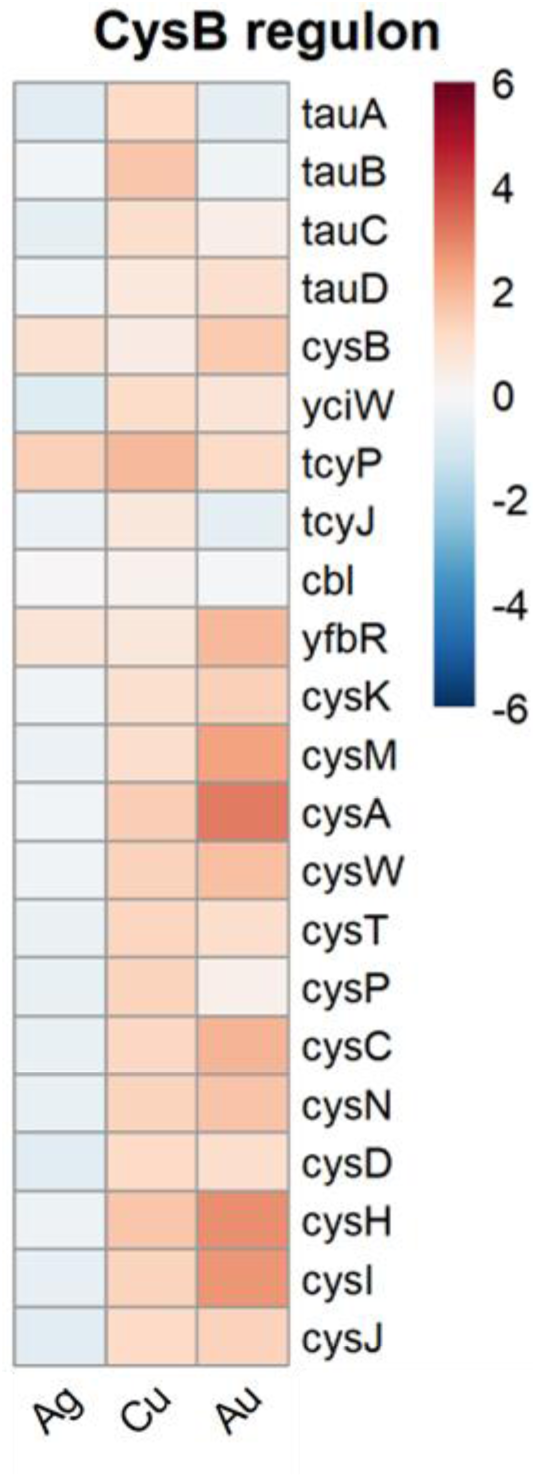
Average log_2_-fold change in gene expression for CysB-regulated genes in *E. coli* K12 BW25113. Each column corresponds to a different experimental condition: silver nitrate (Ag), copper sulfate (Cu), and tetrachloroauric acid (Au). Every experiment was contrasted against a non-metal challenge control, with three biological trials each.

While Cu and Au reported high expression activities in the CysB regulon, Ag did not elicit the same type of response, having most of this regulon down-regulated. Despite this contrast, the CysB-regulated *tcyP* appeared as an UP DEG across our three metal treatments. This gene codes for a L-cystine/L-cysteine shuttle transport system, which provides reducing equivalents to the periplasm, helping restore redox balance as part of the *E. coli* response to oxidative stress (68). This gene, along with the CysB regulon, was also significantly up-regulated in a previous acclimation experiment using gallium nitrate (37). These observations suggest that the TcyP exchanger could play an important role in protecting against oxidative stress induced by toxic metals over longer exposure times.

Our Au experiment yielded the transcriptional profile that differed the most against its non-metal treated control, with 46.89% of our strain’s coding sequences being differentially expressed. Part of this contrast can be explained by what seems to be an increase in protein synthesis. The [Translation iModulon](https://imodulondb.org/iModulon.html?organism=e_coli&dataset=precise1k&k=1) (**Figure 5**) presents the highest activity in our Au treatment. This cluster of co-expressed genes is mostly composed of ribosomal subunit proteins related to translation or ribosomal structure and biogenesis (27). One of the common transcription factors of 44 out of 53 genes from this group is DksA, an autoregulated DOWN DEG (log_2_-fold change: −1.12) which controls the expression of a variety of operons related to amino acid biosynthesis and ribosomal subunits. Furthermore, the up-regulation of several amino acid biosynthesis regulons (MetJ, TrpR, ArgP, ArgR, TyrR; **Table 4**) and iModulons (Histidine, Tyr/Trp/Phe, Tryptophan, Leu/Val/Ile, Cysteine-2; **Figure 4**) suggests an increased demand of amino acids for translation. This evidence leads us to believe that Au exposure is somehow leading to protein breakdown or ribosome disassembly at a greater degree than Ag and Cu, eliciting this kind of response. While direct evidence of gold ion toxicity targeting ribosomes is limited, there is precedent of gold nanoparticles affecting ribosome-tRNA binding by another study (69). It is reasonable to assume a possible path of toxicity by means of interactions between gold ions with exposed thiol and thioether groups from ribosomal proteins, similar to previous observations with auranofin (70).

Another regulon that stood out from our Au experiment is the one controlled by the <u>z</u>inc <u>u</u>ptake regulator Zur. This transcription factor uses four Zn(II) atoms in its active DNA-binding form, repressing the transcription of its target genes (71) including the zinc uptake system ZnuABC and its chaperone ZinT. All of Zur’s target genes are up-regulated in our Au condition, with only one just below significance levels (**Figure 12**). This observation is consistent with the down-regulation of z*ntA*, which encodes for a P-type ATPase that effluxes Zn(II) ions from the cytoplasm to the periplasm(72). *zntA* transcription is activated by ZntR in the presence of Zn(II), and its activation is also dependent on the availability of these ions. Therefore, this could be an indication that our Au treatment may be leading into a zinc-deprived state, commanding a higher demand of Zn(II).

**Figure 12.**
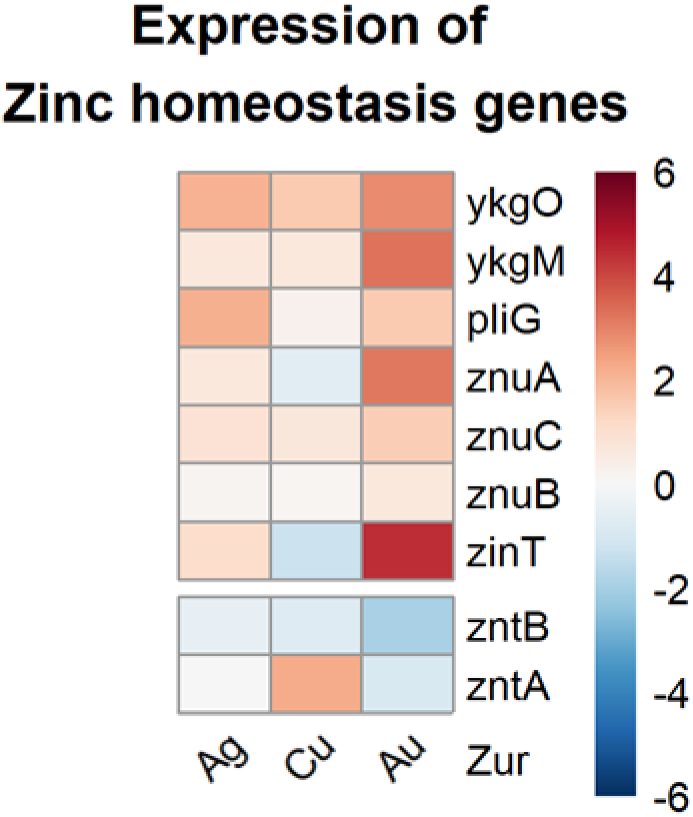
Average log_2_-fold change in gene expression for genes that are known to play a central role in maintaining zinc homeostasis in *E. coli* K12 BW25113. The first group of genes are regulated by Zur, while the lower pair are regulated by ZntR. Each column corresponds to a different experimental condition: silver nitrate (Ag), copper sulfate (Cu), and tetrachloroauric acid (Au). Every experiment was contrasted against a non-metal challenge control, with three biological trials each.

### Concluding remarks

The longstanding dogma for metal toxicity in bacteria has consisted of increased oxidative stress that ultimately results in bacterial death. This hypothesis is rooted in evidence, as multiple studies have correlated the production of ROS in bacteria when exposed to metals. However, this has led to the misconception that all metals behave the same way, and thus bacteria respond to all of them similarly. The observations stemming from our study add layers to this premise. We have provided a model that depicts shared and unique traits in the *E. coli* adaptive and intrinsic gene expression response to each antimicrobial coinage metal at sublethal concentrations. Crucially, we interrogate bacterial physiology over a prolonged exposure where the culture has already acclimated to metal stress, giving clues into what type of response is needed for sustained growth. Disruption in iron metabolism is to be expected as a shared response since [Fe-S] centers are prime targets for free radicals, promoting activity in the Fur regulon. Activation of the copper homeostasis system Cus and Cue by Cu was also expected, although it is interesting that Au only activated the latter while both were relatively unresponsive to Ag. Induction of the Cpx response is also consistent with previous observations of Ag toxicity attacking the cell envelope. The stark difference in transcriptomic profile elicited by Au was a surprise, majorly characterized by increased gene expression in sulfur metabolism pathways (controlled by CysB, also elicited by Cu), translation and ribosome biogenesis genes, and the zinc uptake regulon Zur (**Figure 7**). These observations fill knowledge gaps of metal-bacteria interactions, going beyond the acute toxicity models to gain a better understanding of bacterial physiology adapted to grow in the presence of these antimicrobial metals.

## METHODS

### Sublethal Concentration Determination

*E. coli* K12 BW25113 stored at −70 °C was thawed and cultured in Lysogeny Broth (LB) media (10 gr/L tryptone, 10 gr/L NaCl, 5 gr/L yeast extract) overnight at 37 °C 150 rpm in a shaker incubator to recover the bacterial strain. For each experiment in a 250 mL Erlenmeyer flask, 500 µL of a bacterial overnight culture were used to inoculate 7.5 mL of 2X M9-glucose minimal media diluted to a 1X concentration (6.8 g/L Na_2_HPO_4_, 3 g/L KH_2_PO_4_, 1 g/L NH_4_Cl, 0.5 g/L NaCl, 4 mg/L glucose, 0.5 mg/L MgSO_4_, and 0.1 mg/L CaCl_2_) with 7.5 mL of metal salt (AgNO_3_, CuSO_4_, or HAuCl_4_) in aqueous solution. Untreated positive controls for bacterial growth used sterile deionized water ((dd)H_2_O) instead of the metal salt component, while negative controls for media sterility had culture media and (dd)H_2_O but no inoculum.

Each flask was incubated in a shaker incubator at 37 °C 150 rpm. Optical density at 600nm (OD600) was determined using a spectrophotometer. Relative to the untreated control, possible sublethal inhibitory concentrations were determined to have either a) an extended lag phase that suggest an adjustment of the bacterial physiology to the metal-spiked media, b) a less steep slope during the exponential growth phase indicating reduced doubling times, or c) a reduced OD600 value (between 45% and 95%) compared to the OD600 from the untreated control after 10 hours of growth defining inability to reach saturating cell numbers.

### Total RNA Sample Preparation

Experimental treatments of AgNO_3_ 7 µM, CuSO_4_ 39 µM, or HAuCl_4_ 10 µM were set up along with their respective non-metal controls as described above. Ten hours after inoculation, 2 mL of bacterial culture were centrifuged at 14,000 X g for 2 min to pellet the cells. Total RNA was then extracted using RiboPure-Bacteria RNA Isolation Kit AM1925 (Invitrogen, USA) according to the manufacturer’s recommendations, including DNase treatment. Three biological trials were set up for both conditions.

### RNA Sequencing

Samples were sent to the Centre for Health Genomics and Informatics and UCDNA Sequencing facility (Cumming School of Medicine, University of Calgary, Canada) for sequencing. Ribosomal RNA depletion was performed using the NEB rRNA Depletion Bacterial Module (New England Biolabs, USA). The cDNA library preparation for Illumina sequencing was executed using the NEBNext Ultra II Directional RNA Library Prep Kit and NEB Indexing Kit (New England Biolabs, USA). After adaptor ligation and amplification, the average library fragment size (in base pairs) for each dataset was as follows: Ag 371, Cu 366, and Au 362. Libraries were quantified using KAPA qPCR Library Quantification Kit for Illumina platforms, then pooled and loaded on the Illumina MiSeq sequencer at a concentration of 12 pM. The libraries were sequenced using a 2×75bp 150 cycle run, with a total output of 25M read pairs.

### Differentially Expressed Genes (DEGs) Identification, Quantification and Analysis

Sequencing files quality control, quantification of gene expression and DEG analysis was performed in a variety of R-studio library scripts. Initial FASTQ files (submitted to NCBI Sequence Read Archive under BioProject accession numbers PRJNA1256886 – Copper, PRJNA1256887 – Silver, PRJNA1256888 – Gold) were assessed and trimmed with the Bioconductor package *rfastp* (73) (version 1.10). Gene expression was assessed by aligning to the *E. coli* K12 BW25113 RefSeq assembly annotation file GCF_000750555.1 using the Bioconductor package *Rsubread* (74) (version 2.14.2), with counts assigned to the CDS meta-feature (**Tables S1-S3**). Finally, the counts matrix for the 4,378 identified features was imported into the Bioconductor package *DESeq2* (75) (version 1.40.2) for DEG determination. P-values were calculated using the Wald test. The adjusted p-value (p-adj) for multiple test adjustment (Benjamini-Hochberg correction, FDR < 0.05) was set to 0.05. DEGs were defined as those with a p-adj value < 0.05 and whose expression had an absolute fold-change (FC) of at least 2 (|log_2_ FC| > 1) when contrasting each metal salt treatment versus their respective unchallenged controls; those with a negative FC were deemed as down-regulated genes in each metal salt treatment, while a positive FC indicated up-regulation (**Tables S1-S3**). Functional enrichment of individual genes was further annotated through the UniProt and EcoCyc databases. Functional network enrichments were performed by submitting lists of genes of interest to the STRING web software (34).

The ISMARA web software (Swiss Institute of Bioinformatics, Basel, Switzerland) (26) was used to detect motifs in the promoters of the *E. coli* K12 genome and infer the activity of gene regulators (including transcription factors, small RNA and RNA-polymerase subunits) based on the expression of their target genes. Our FASTQ files were uploaded to the ISMARA web server and averaged together by treatment.

For our co-expression analysis, the *E. coli* PRECISE-1K dataset (28) was utilized, and the activity of each of the 201 iModulons was determined for both conditions of our experiment as indicated by Lamoreux *et al*. The difference in activity between both conditions represents differential iModulon activity between our treatments, with positive values indicating higher activity in each metal salt treatment, and negative ones representing lower activity. Statistical significance was calculated using the cumulative log-normal distribution of each iModulon across all the conditions.

## DATA AVAILABILITY

Processed transcriptomic data (sequencing files, raw counts matrix and *DESeq2* output table) are available under NCBI Gene Expression Omnibus accession numbers GSE295970 (copper, reviewer token **knqzugoelzovbmd**), GSE295971 (gold, reviewer token **ynuboesspxezbgb**), and GSE295972 (silver, reviewer token mlmbkmcolbwpbet).

## ACKNOWLEDGMENTS

We would like to acknowledge the Centre for Health Genomics and Informatics and UCDNA Sequencing facility (Cumming School of Medicine, University of Calgary, Canada) for their help with RNA-seq library preparation and sequencing. DASA was supported by an Eyes-High Doctoral Scholarship from the University of Calgary. This work was funded by a Discovery grant awarded to RJT from the Natural Sciences and Engineering Research Council of Canada (RGPIN 2020-03877).

## Author contributions were as follows

conceived and designed the study— D.A.S.A. and R.J.T.; practical performance—D.A.S.A. and A.M.; processed the data— D.A.S.A.; wrote the first draft of the paper—D.A.S.A and R.J.T.; and participated in data analysis and manuscript editing—D.A.S.A., A.M., and R.J.T. All authors have read and agreed to the published version of the manuscript. We declare there are no competing financial interests.

